# Elucidating a locus coeruleus-hippocampal dopamine pathway for operant reinforcement

**DOI:** 10.1101/2022.10.13.512041

**Authors:** Elijah A. Petter, Isabella P. Fallon, Ryan N. Hughes, Glenn D. R. Watson, Warren H. Meck, Francesco Paolo Ulloa Severino, Henry H. Yin

## Abstract

Animals can learn to repeat behaviors to earn desired rewards, a process commonly known as reinforcement learning. While previous work has implicated the ascending dopaminergic projections to the basal ganglia in reinforcement learning, little is known about the role of the hippocampus. Here we report that a specific population of hippocampal neurons and their dopaminergic innervation contribute to operant self-stimulation. These neurons are located in the dentate gyrus, receive dopaminergic projections from the locus coeruleus, and express D1 dopamine receptors. Activation of D1+ dentate neurons is sufficient for self-stimulation: mice will press a lever to earn optogenetic activation of these neurons. A similar effect is also observed with selective activation of the locus coeruleus projections to the dentate gyrus, and blocked by D1 receptor antagonism. Calcium imaging of D1+ dentate neurons revealed significant activity at the time of action selection, but not during passive reward delivery. These results reveal the role of dopaminergic innervation of the hippocampus in supporting operant reinforcement.

## Introduction

In operant learning, animals modify their action repertoires to earn desired rewards. Previous work on the neural substrates of such learning has focused on the striatum and the midbrain dopaminergic projections that target the striatum (Wise, 2004; Yin et al., 2005; Kravitz et al., 2013; Rossi et al., 2013; Yttri and Dudman, 2016). This type of learning is often thought to be distinct from declarative or episodic learning, which require the hippocampus and medial temporal lobe structures (Mishkin, 1984; Morris et al., 1986; Milner et al., 1998; Eldridge et al., 2000). On the other hand, work in both humans and rodents has also implicated the hippocampus in reward processing and motivated behavior, though the underlying mechanisms remain unclear (Adcock et al., 2006; Gauthier and Tank, 2018).

Midbrain dopamine neurons have been implicated in reinforcement learning (Schultz et al., 1997; Tsai et al., 2009; Rossi et al., 2013). The hippocampus is also a target of dopaminergic projections. Dopamine receptors are expressed in the hippocampus, and in mice D1-class receptor expression is common in the dentate gyrus (DG) region (Gangarossa et al., 2012; Kempadoo et al., 2016). However, these dopaminergic projections come from locus coeruleus (LC) (Kempadoo et al., 2016), rather than the major dopamine cell groups in the ventral tegmental area (VTA) and substantia nigra pars compacta (SNc), which supply dopamine to the basal ganglia (Björklund and Dunnett, 2007; Ikemoto, 2007).

In this study we examined the contribution of dopaminergic signaling in the DG to operant learning and behavior. We found that mice could learn to perform a new action (pressing a lever) for optogenetic activation of D1+ neurons in the DG. In addition, using both optogenetics and *in vivo* pharmacological manipulations, we found that activation of LC dopaminergic neurons that project to the DG can also produce self-stimulation, and this effect depends on the activation of D1-like receptors. Finally, using *in vivo* calcium imaging, we found that D1+ DG neurons were more related to the goal-directed actions than simply passive reward, and play a prominent role in operant behavior.

## Results

To understand the role of D1+ neurons in the hippocampus, we tested whether selective stimulation of these neurons can reinforce operant behavior using a self-stimulation paradigm. We injected either a Cre-dependent channelrhodopsin (AAV5-DIO-ChR2) or a fluorescent control (DIO-eYFP) into D1-Cre mice (D1::ChR2^DG^ or D1::eYFP^DG^), producing selective expression of the excitatory opsin in D1+ neurons in the dentate gyrus (**Figure 1A-B)**. Mice received photostimulation (500 ms, 20 Hz, 15 ms pulse width) following lever pressing on a fixed ratio schedule of reinforcement (**Figure 1C**). All D1::ChR2^DG^ mice learned to press a lever for stimulation, whereas control mice did not (**Figure 1D**). These results suggest that D1::ChR2^DG^ stimulation is sufficient to reinforce lever pressing. Interestingly, this form of self-stimulation is remarkably resistant to extinction, persisting after 8 days without any photostimulation.

**Figure 1.**
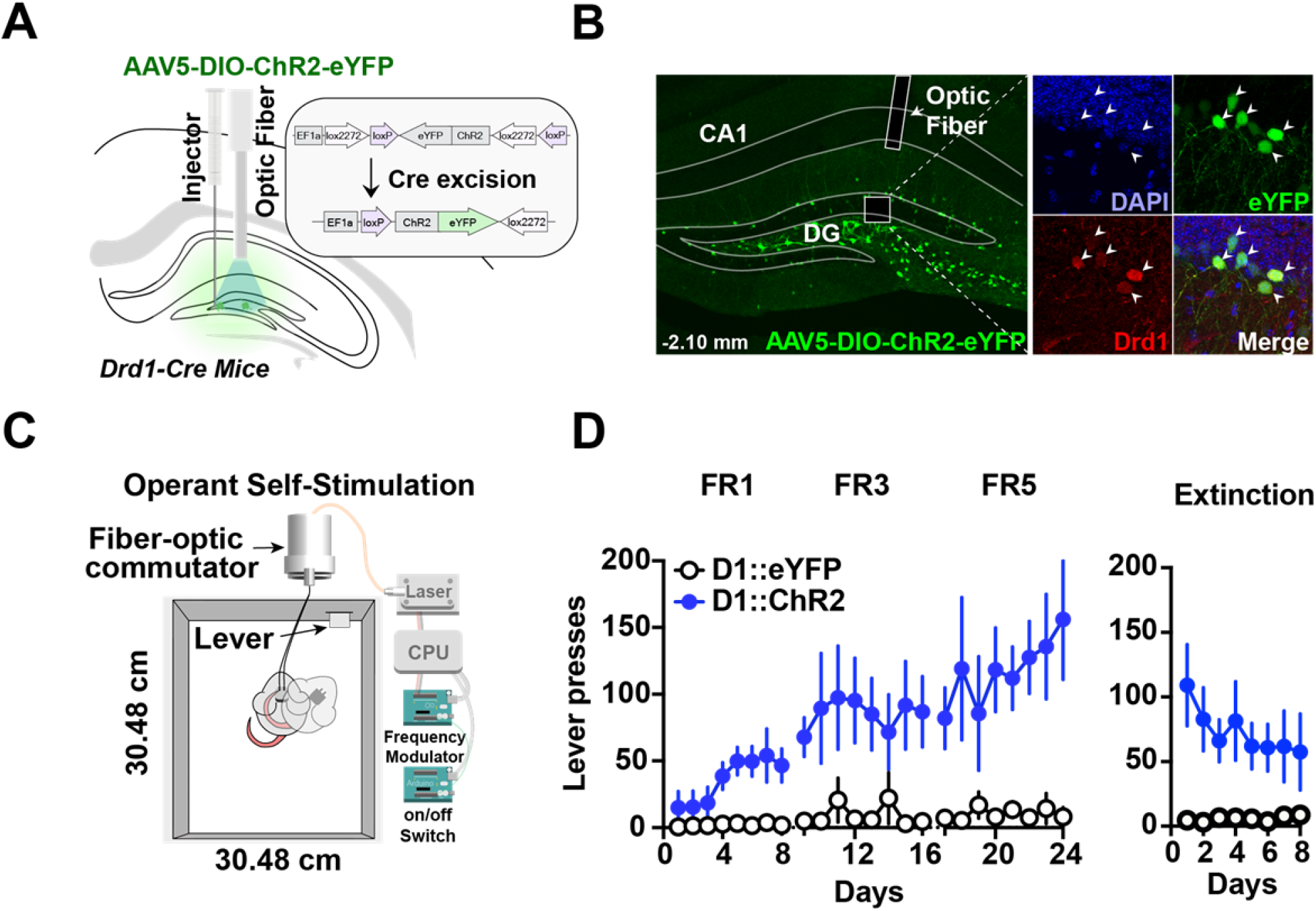
Optogenetic stimulation of D1+ neurons in the dentate gyrus is sufficient for operant self-stimulation. (**A**) Schematic of optic fiber placement above virally infected D1+ dentate gyrus (DG) neurons in *Drd1-Cre* transgenic mice. (**B**) *Left*, coronal section showing ChR2 expression in the DG. *Right*, magnified view of AAV infected DG neurons from inset colocalized with D1 receptors. White arrows indicate cell bodies. (**C**) Schematic of the operant self-stimulation chamber. (**D**) Lever pressing rate across three fixed ratio (FR1, FR3, FR5) schedules of reinforcement, and extinction (8 days each) for D1:Chr2-DG animals (*N =* 8) and eYFP (*N =* 5) controls. D1::Chr2-DG mice self-stimulated significantly more than controls (Two-way RM ANOVA, Group [ChR2 or eYFP] x Day, main effect of group *F(1,11)* =9.474, *p* = 0.01, main effect of Day, *F(23, 253)* = 1.93, *p* = 0.0078; no significant interaction *F(23,253)* = 1.418, *p* = 0.10). During extinction, there was a significant main effect of group: *F(1,88)* =33.79, *p* < 0.0001, no significant effect of Day: *F(7, 88)* = 0.26, *p* = 0.966, and no interaction: *F(7, 88)* =0.327, *p* = 0.940). Means +/− SEM for all graphs. DG, dentate gyrus; LC, Locus Coeruleus; scp, superior cerebellar peduncle; DAPI, 4′,6-diamidino-2-phenylindole. **** *p* < 0.0001

Next, using retrograde tracing methods, we were able to map projections to the DG (**Figures 2A-D & Supplementary Figure 1**). We confirmed significant LC projections to the DG, but we did not find significant VTA or SNc projections (**Figure 2E, H & Table 1**). Retrograde labeling of DG-projecting LC neurons is colocalized with tyrosine hydroxylase (TH), a marker for catecholamine neurons (e.g. dopamine, norepinephrine; **Figures 2F-H**). In contrast, there was no labeling in the VTA (**Figure 2H & Supplementary Figure 1**). This finding suggests that the DG receives TH+ projections from the LC rather than VTA.

**Table 1:** Colocalization of tyrosine hydroxylase with retrograde DG labeling in the Locus Coeruleus. Colocalization of TH and Retro-cre labeling (N=8; 4 mice, 2 hemispheres), showing that at least some of the LC-DG neurons are TH positive. In contrast, no colocalization was found with retro-Cre and TH labeling in the VTA (N=6; 3 mice, 2 hemispheres), VTA slices were not taken from one animal.

| Animal | Retro-cre label VTA (from DG) | Retro-Cre label LC (from DG) | LC colocalized with TH |
| --- | --- | --- | --- |
| Mouse 1 (LH) | 0 | 18 | 4 |
| Mouse 1 (RH) | 0 | 32 | 7 |
| Mouse 2 (LH) | 0 | 5 | 2 |
| Mouse 2 (RH) | 0 | 13 | 6 |
| Mouse 3 (LH) | 0 | 20 | 12 |
| Mouse 3 (RH) | 0 | 31 | 15 |
| Mouse 4 (LH) | n/a | 22 | 12 |
| Mouse 4 (RH) | n/a | 14 | 5 |

**Figure 2:**
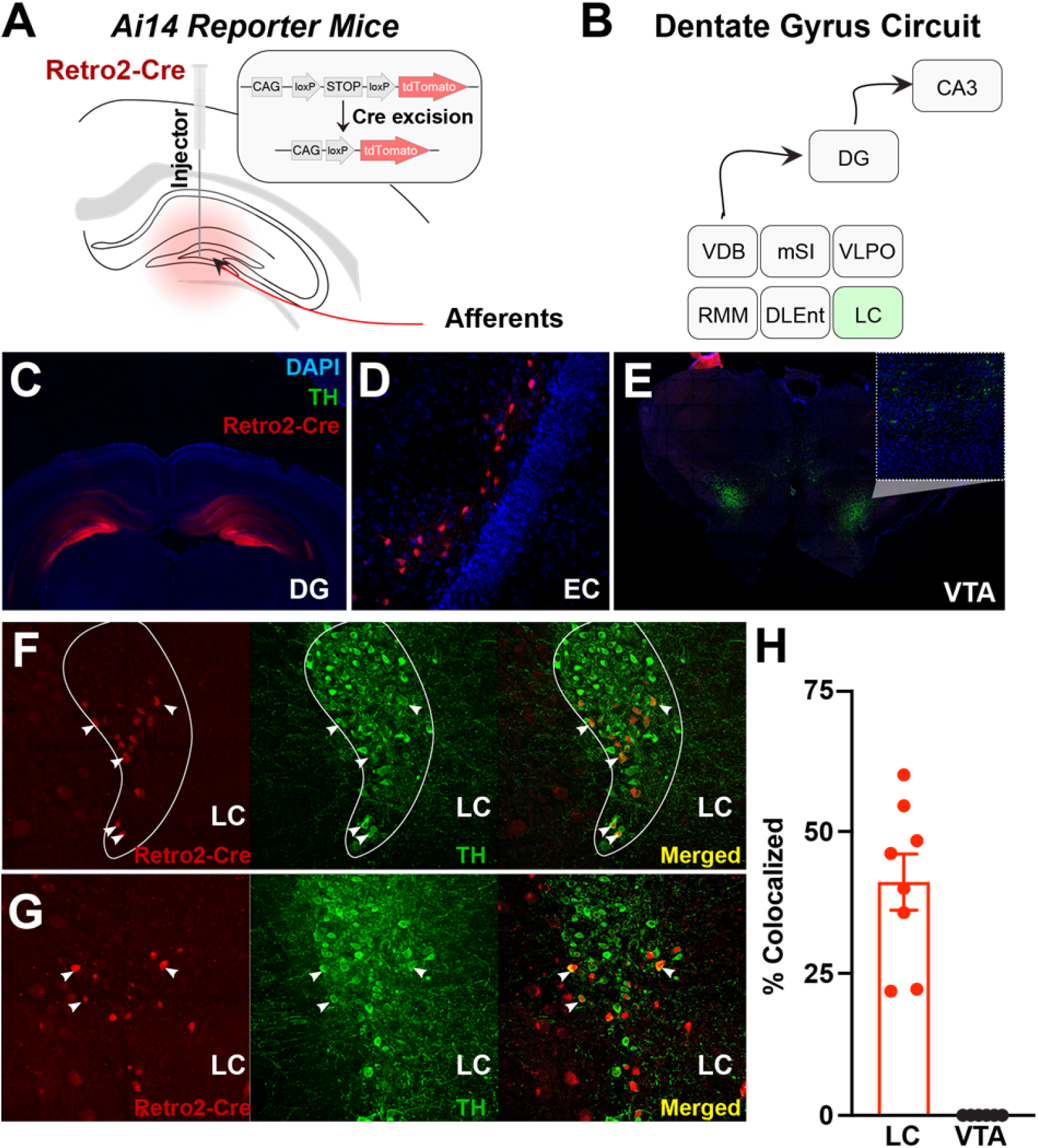
Retro-Cre tracing shows that the main catecholamine input to the dentate gyrus is the Locus Coeruleus, not ventral tegmental area. (**A**) Schematic of Retro2-Cre injection into the dentate gyrus in Ai-14 reporter mice. (**B**) Schematic summarizing all brain regions that project to the DG. Only LC is TH+. Abbreviations – Vertical diagonal band (VDB), Medial septal nucleus (MSN), Ventral lateral preoptic area (VLPO), retro mammillary bodies (RMM), dorsal lateral entorhinal cortex (DLEnt) (**C**) Injection site of the Retro2 showing the Cre-positive neurons.(**D-G**) Retrograde labeling of neurons in canonical brain regions that project to the hippocampus. (**D**) entorhinal cortex (EC). (**E**) Limited retrograde labeling of neurons in the VTA, colocalized with tyrosine hydroxylase (TH). (**F & G**) Retrograde labeling of LC neurons in two out of four mice, colocalized with tyrosine hydroxylase. (**H**) Percent of colocalized neurons in the LC (*N =* 8; 4 mice x 2 hemispheres) and VTA (*N =* 6; 3 mice x 2 hemispheres). Unpaired t test, p < 0.0001. Mean and +/- sem.

We then tested whether the LC-DG projection is responsible for the self-stimulation effect observed. In order to manipulate the LC-DG pathway selectively, we injected *AAV-Retro2-Cre* into the DG and a Cre-dependent ChR2 (*AAV5-DIO-ChR2)* into the LC (**Figure 3A-B**). We found that ChR2^DG-LC^ (*N = 6*) mice also showed self-stimulation that is comparable to the stimulation of D1::ChR2^DG^ neurons (**Figure 3C**).

**Figure 3.**
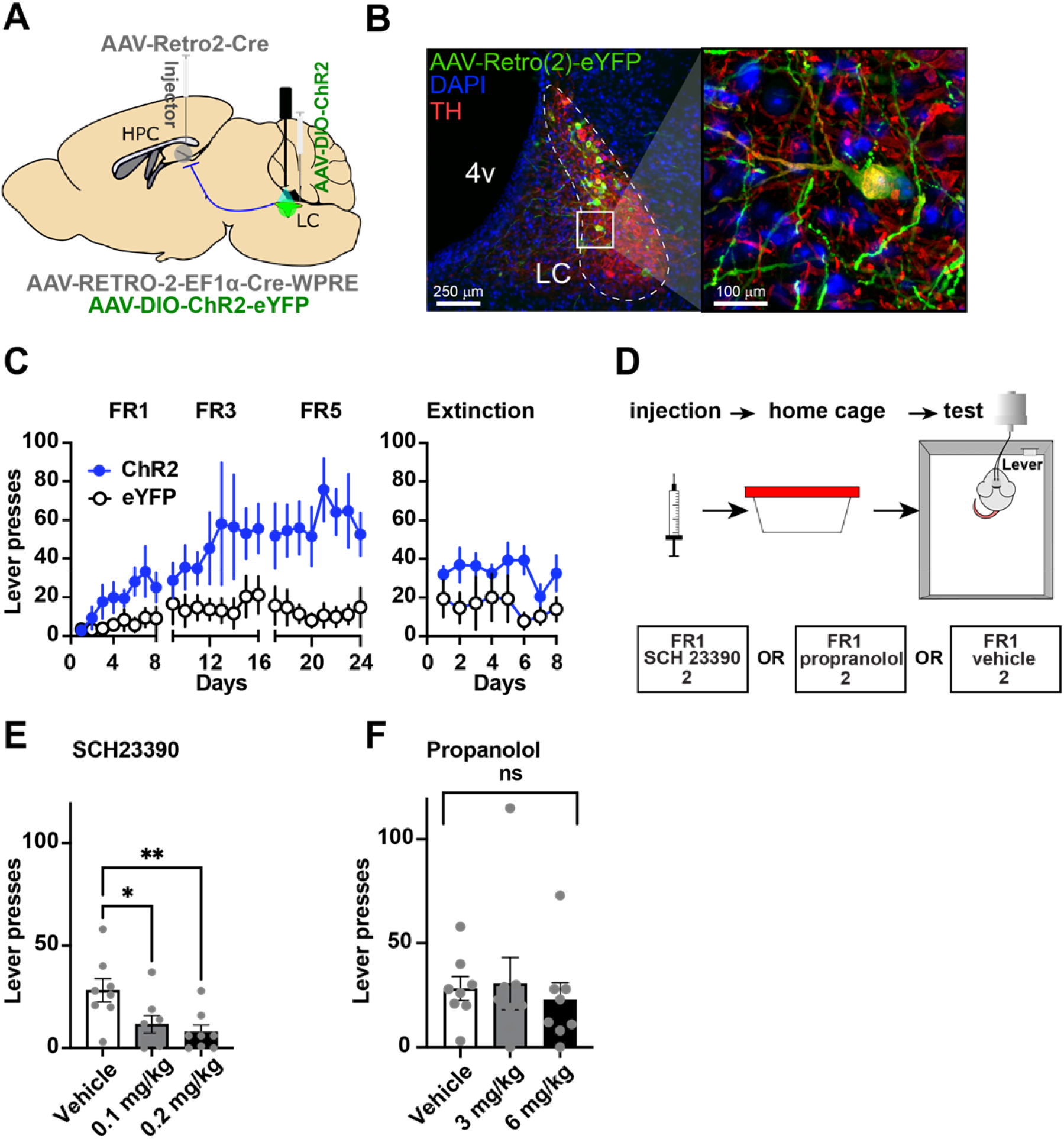
Locus Coeruleus projections to the dentate gyrus contribute to operant learning. **(A)** Schematic showing selective targeting of locus coeruleus (LC) projections to the dentate gyrus (DG). Cre expression was induced by first injecting AAV-Retro2-Cre into the DG. A Cre-dependent virus (AAV5-DIO-ChR2-eYFP) was then injected into the LC before optic fiber implantation. (**B**) Left: Representative coronal section showing Cre-dependent ChR2 expression in the LC. Right: Magnified view from inset showing eYFP colocalization with tyrosine hydroxylase (TH) LC neurons. (**C**) Lever presses per session for three fixed ratio (FR1, FR3, FR5) schedules of reinforcement, and extinction for Retro::ChR2 DG-LC animals (N = 6) and eYFP (N = 5) controls. Mice with ChR2 expressed in LC self-stimulated significantly more than controls (Two-way RM ANOVA Group [ChR2 or eYFP] x Day, significant effect of group: F(1,10) = 6.59, p = 0.028; significant effect of day, F(23,230) = 2.564, p = 0.0002; no interaction: F(23,230) = 1.443, p = 0.09. (**D**) Left, design of pharmacological experiments. After FR1 training, mice received IP injections of antagonists for either D1 (SCH 23390), or NE-beta receptors (propranolol). **(E)** D1-antagonist SCH 23390 significantly reduced self-stimulation of DG-projecting LC neurons: F(2, 14) = 6.9, p = 0.008. Post hoc analysis (Dunnett’s) shows that both doses reduced lever pressing relative to controls (0.1 mg/kg, p = 0.002; 0.2 mg/kg, p = 0.007). **(F)** The NE antagonist propranolol did not have any effect: F(2, 14) = 0.290, p = 0.753. Means +/− SEM. HPC, hippocampus; 4v, fourth ventricle. * p<0.05, **p <0.01.

The LC-DG projection releases both norepinephrine and dopamine(Kempadoo et al., 2016; Takeuchi et al., 2016). It is unclear which transmitter is responsible for the self-stimulation effect, though our observation on D1+ DG neurons (**Figure 1**) suggest that dopamine might be responsible. Consequently, to determine which of these transmitters is responsible for the observed effects, we used pharmacological manipulations in combination with pathway-specific optogenetic manipulations using the same self-stimulation paradigm (**Figure 3E & Figure 4**). Mice (*N = 8*) trained on self-stimulation were tested after either receiving systemic injections of a β-adrenoceptor antagonist (propranolol), or a D1-antagonist (SCH 23390). β-adrenoceptor blockade did not produce any significant effects (**Figure 3D, F)**. In contrast, D1-antagonist significantly impaired self-stimulation (**Figure 3E**).

**Figure 4.**
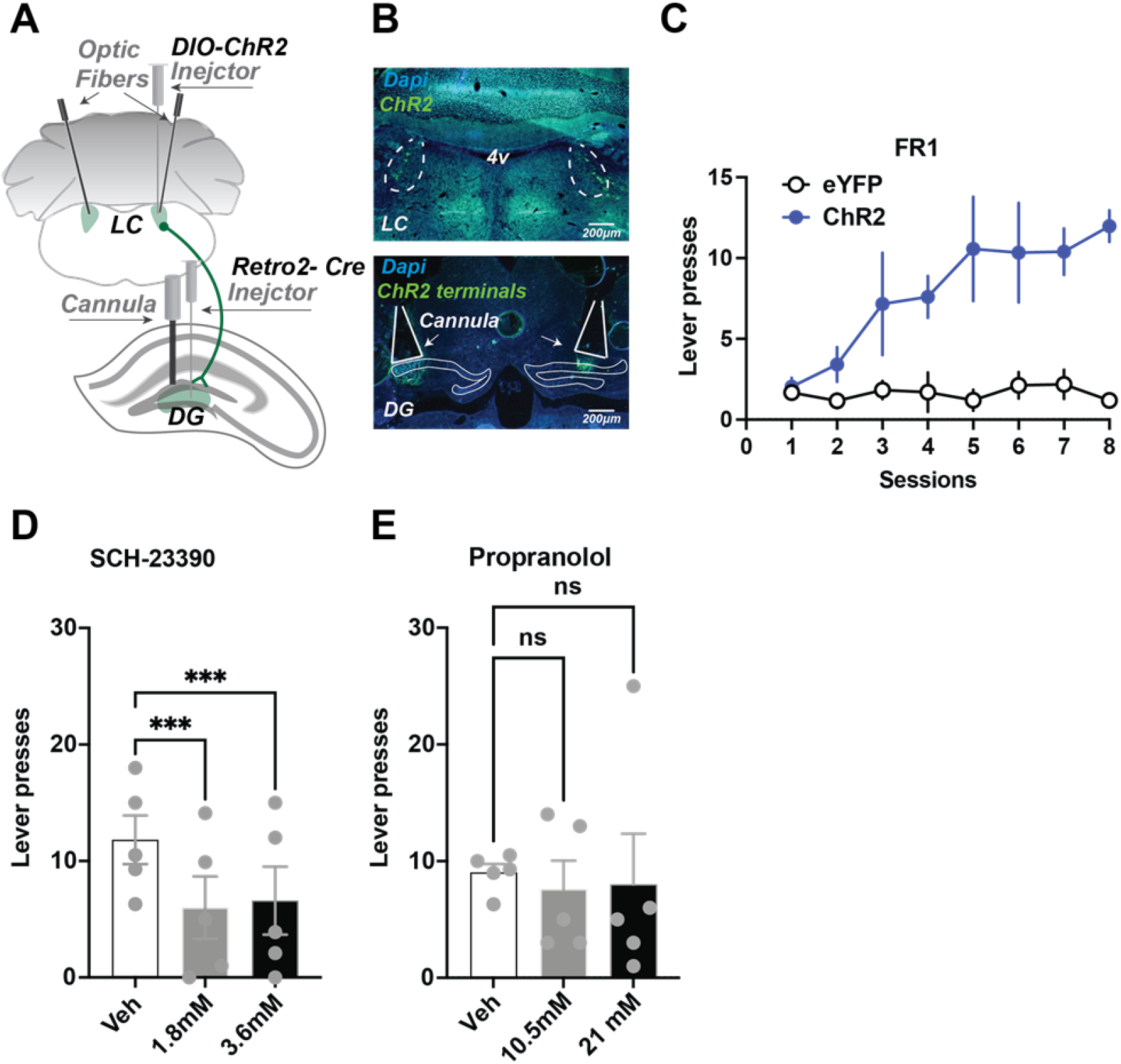
Local infusions of D1 but not NE beta antagonist into the DG reduces self-stimulation. **(A)** Schematic showing injection strategy for local drug infusions into the DG during self stimulation of locus coeruleus (LC) neurons that project to the dentate gyrus (DG) (N=5). Cre expression was induced in LC neurons projecting to the DG by first injecting AAV-Retro-2 into the DG. An injection of a Cre-dependent virus (AAV5-DIO-ChR2-eYFP) was then made in the LC before optic fiber implantation. Canulae were used to inject DA and NE antagonists into the DG. **(B)** Top, representative coronal section showing ChR2 expression in the LC. Bottom, ChR2 terminals in DG and cannula tracks. **(C)** Acquisition of lever pressing Retro::ChR2 DG-LC mice or Retro::eYFP DG-LC (controls). Experimental animals self-stimulated more than controls (Two-way ANOVA [Day x Group], effect of Day F(7, 84) = 5.222, p < 0.0001; effect of group, F(1,12) = 37.98, p < 0.0001; interaction F(7,84) = 4.932, p < 0.0001). **(D)** One-way RM ANOVA showed D1 antagonist SCH-23390 significantly reduced lever pressing. There is a significant drug effect: F(2,8) = 28.22, p = 0.0002. Both doses produced significant suppression of lever pressing (1.8 mM, p =0 .0003; 3.6 mM, p = 0.0005). **(E)** NE antagonist propranolol had no significant effect on self stimulation. F(2,8) = 0.05, p = 0.951. Means +/− SEM for all graphs. DG, dentate gyrus; LC, locus coeruleus; 4v, fourth ventricle.

To activate DG-projecting LC neurons robustly, we targeted the cell-bodies of the DG-LC projections. But the LC has broad projections to many brain areas (Schwarz and Luo, 2015). To verify that our self-stimulation effects were not due to activation of LC collaterals in other regions, we performed local infusions of antagonists (**Figure 4A-C**, *N = 5*). Infusions of a D1-antagonist into the DG significantly impaired self-stimulation, whereas propranolol showed no significant group differences on self-stimulation (**Figure 4D-E**). These results suggest that the reinforcing effects of LC-DG stimulation is due to the activation of D1 receptors by dopamine, rather than by norepinephrine.

Based on our self-stimulation results, we hypothesized that DG D1+ neurons may be preferentially activated during operant conditioning in general, including during actions that result in natural rewards rather than simply optogenetic stimulation of the LC-DG pathway. To test this, we performed *in vivo* calcium imaging of DG D1+ neurons while performing an operant lever pressing task with food reward. We implanted a gradient index lens above the DG in D1-Cre mice (*N = 5*) injected them with a Cre-dependent calcium indicator (*AAV9-syn-FLEX-jGCamp7f*) (**Figure 5A-B**).

**Figure 5.**
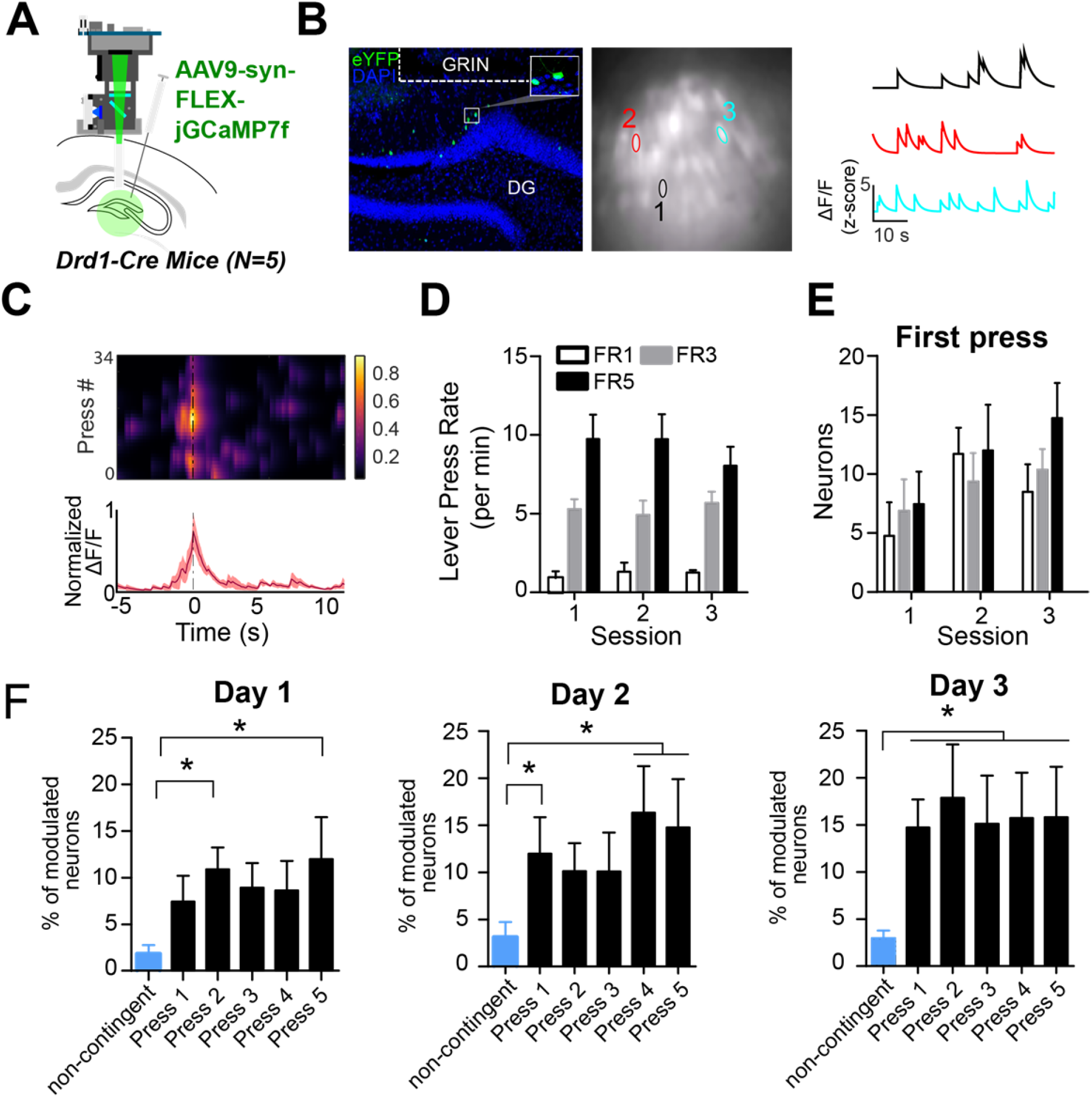
D1+ dentate gyrus neurons are significantly modulated by lever pressing during operant conditioning. (**A**) A schematic of a UCLA miniscope and GRIN lens implanted over jGCamp7f-infected D1+ cells in the dentate gyrus (DG) of *Drd1-Cre* mice for *in vivo* imaging (N=5). (**B**) *Left:* Representative coronal section showing GCaMP7f expression in D1 neurons of the DG. GRIN lens, marked in white. *Middle*: Imaging field of view with contours of identified neurons. *Right*: example calcium traces significantly modulated by lever presses. (**C**) An example neuron showing increased calcium transient during lever pressing. *Top*: Heat map shows normalized calcium activity aligned to a single press as a function of time during fixed-ratio (FR) 5 trials. *Bottom*: Averaged calcium activity across all presses. (**D**)The press rates for FR1, FR3, and FR5 for each day of testing. The press rates increase across FR schedule. (**E**) The percent of neurons modulated by the first press in each FR schedule across days. Two-way RM ANOVA FR schedule x Day, no effect of FR schedule *F(2,4)* = 0.6702, *p* = 0.5298; effect of day, *F(2,4)* = 4.536, *p* = 0.0213; no effect of interaction *F(2,4)* = 0.6403, *p* = 0.6389 (**F)** The percent of neurons modulated by a non-contingent reward task, or by each press in an FR5 task. Day 1, One-way RM ANOVA *F(4,5)* = 3.389, *p* = 0.0222. Day 2, One-way RM ANOVA *F(4,5)* = 16.19, *p* = 0.0046. Day 3, One-way RM ANOVA *F(4,5)* = 4.807, *p* = 0.0048. Dunnett’s multiple comparison test was used to compare the percent of press modulated neurons to the non-contingent modulated neurons. Significance values are marked (*) for p<0.05.

We then recorded calcium transients from DG D1+ neurons during operant lever pressing for food rewards (**Figure 5A-C**) during lever pressing for food reward on fixed-ratio (FR) schedules (FR1, FR3 and FR5). We found distinct populations of DG D1 + neurons that were modulated by lever pressing. To see if the neural activity is action-contingent, we also used a control task in which pressing is not required. The reward was delivered non-contingently every 20 seconds, preceded by one second of white noise. On this task, there were far fewer significantly modulated DG D1+ neurons (*N = 6*, 3% of total population) compared to the operant task (**Figure 5F, Table 2)**. To verify that the virus targets D1+ neurons in the DG, we quantified the percent of neurons that are virally targeted that express D1 receptors. Using RNA scope, we found that GcAMP-7f was colocalized with D1 receptors (**Figure 6**).

**Table 2:** Calcium imaging neuron counts for each task.

| Task | Animal 1 | Animal 2 | Animal 3 | Animal 4 | Animal 5 |
| --- | --- | --- | --- | --- | --- |
| *Delay Task* | 64 | 62 | 22 | 59 | 100 |
| *FR1 Day 1* | 40 | 53 | 31 | 82 | 85 |
| *FR1 Day 2* | 42 | 48 | 26 | 99 | 45 |
| *FR1 Day 3* | 26 | 48 | 20 | 94 | 46 |
| *FR3 Day 1* | 26 | 54 | 23 | 90 | 32 |
| *FR3 Day 2* | 27 | 71 | 23 | 75 | 33 |
| *FR3 Day 3* | 27 | 75 | 27 | 80 | 33 |
| *FR5 Day 1* | 32 | 91 | 25 | 98 | 43 |
| *FR5 Day 2* | 28 | 106 | 27 | 100 | 57 |
| *FR5 Day 3* | 33 | 106 | 28 | 75 | 57 |
| *FR5 switch Day 1* | 99 | 65 | 18 | 56 | 63 |
| *FR5 switch Day 2* | 100 | 64 | 20 | 49 | 63 |
| *FR5 switch Day 3* | 96 | 66 | 21 | 54 | 62 |
| *Non contingent* | 123 | 94 | 23 | 78 | 96 |

**Figure 6.**
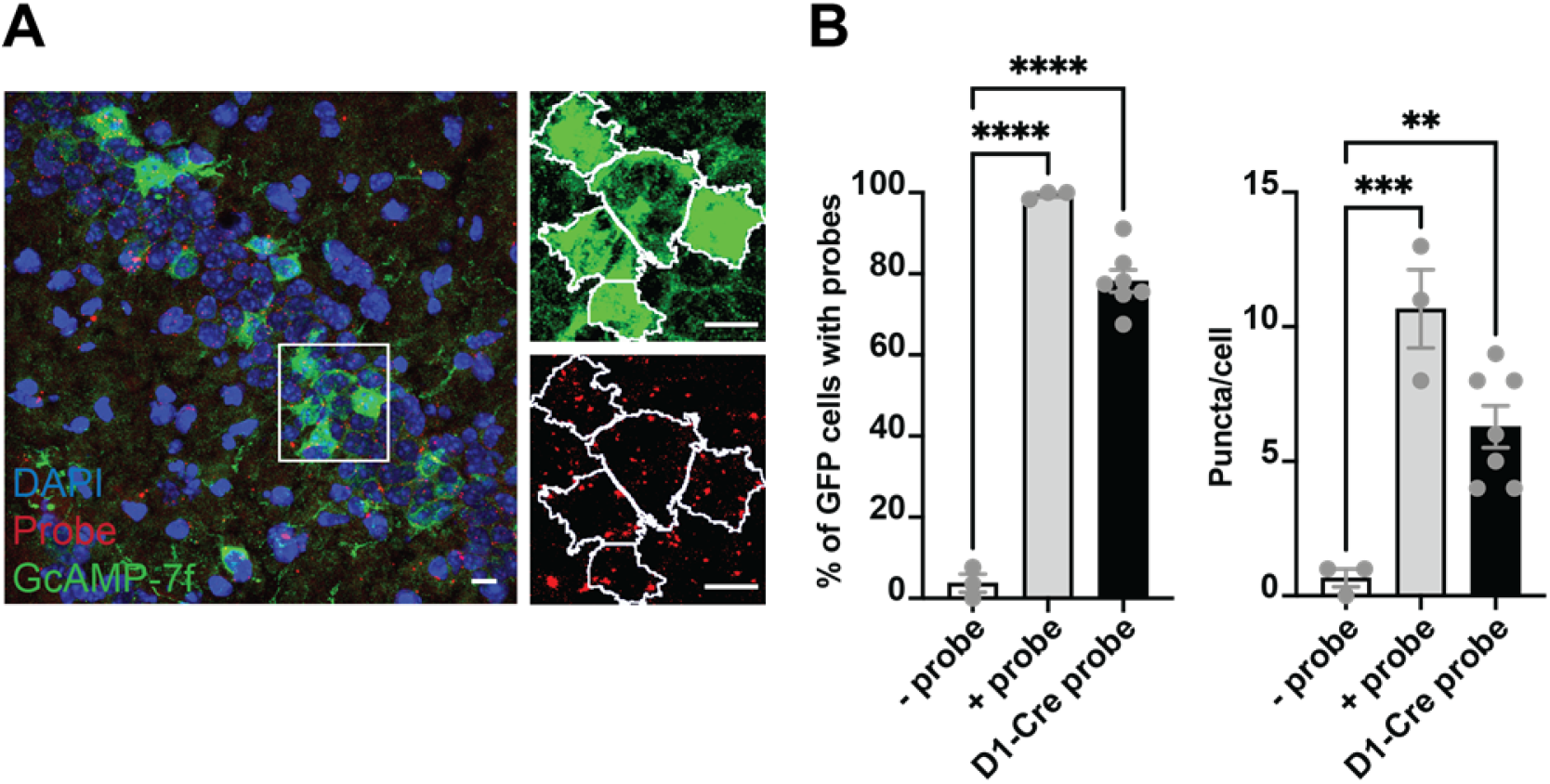
Colocalization of GcAMP-7f with D1 RNA scope probes. **(A)** *Left*, representative coronal section from a Drd1a-cre mouse injected with GCaMP-7f. *Right*, zoomed in view of green (GCaMP7f) and red (probe) channels, showing the outlines (white) of GFP+ cells identified in FIJI. **(B)** *Left*, percent of GFP cells colocalized with the negative control probe (N=3 sections; 1 brain, 9 images), positive control probe (N = 3 sections; 1 brain, 5 images), and Drd1a probe (N = 7 sections; 3 brains, 38 images). One-way ANOVA showed a significant group difference (F = 229.2, p < 0.0001). About 80% of GFP neurons colocalized with the Drd1a, much higher than background negative control probe (Tukey‘s, p < 0.0001). *Right*, there was also a significant group difference in puncta per cell (ANOVA, F = 19.51, p = 0.0004) between Drd1a probe and negative control probe (Tukey‘s, mean diff = -5.619, adjusted p < 0.0052) and positive control probe (Tukey‘s, mean diff = 4.381, adjusted p = 0.0226). Thus the SIO/DIO constructs are only being expressed in D1+ neurons. Scale bars represent 10um.

To determine if the activity of these neurons reflected the spatial locations of the lever pressing or the action of lever pressing itself, we used a discrete trial design with two levers (**Figure 7**). On each trial, one of the two levers was randomly selected to extend into the operant box. Once pressed, the lever would retract. The reward would then be delivered one second later. This task allowed us to compare the neural activity modulated by lever pressing and reward, as well as determine the spatial tuning of the same neurons. We found that several populations of dentate D1+ neurons (*n = 40*, 16.5% of total population) that were significantly modulated at the time of lever pressing (**Figure 7C**). One small population was modulated by reward delivery (*n =14*, 4.9% of total population). Importantly, another population with significantly more neurons were responsive to lever pressing at either lever location (**Figures 7C-D;** *N = 22*, 9.79% of total population). These neurons were not spatially selective, as they were responsive when the lever was presented at different locations. However, we did find a small population that responded to only a single lever (**Figure 7C**; left lever: *n = 15*, right lever: *n = 12*; 4.2 % of total population).

**Figure 7.**
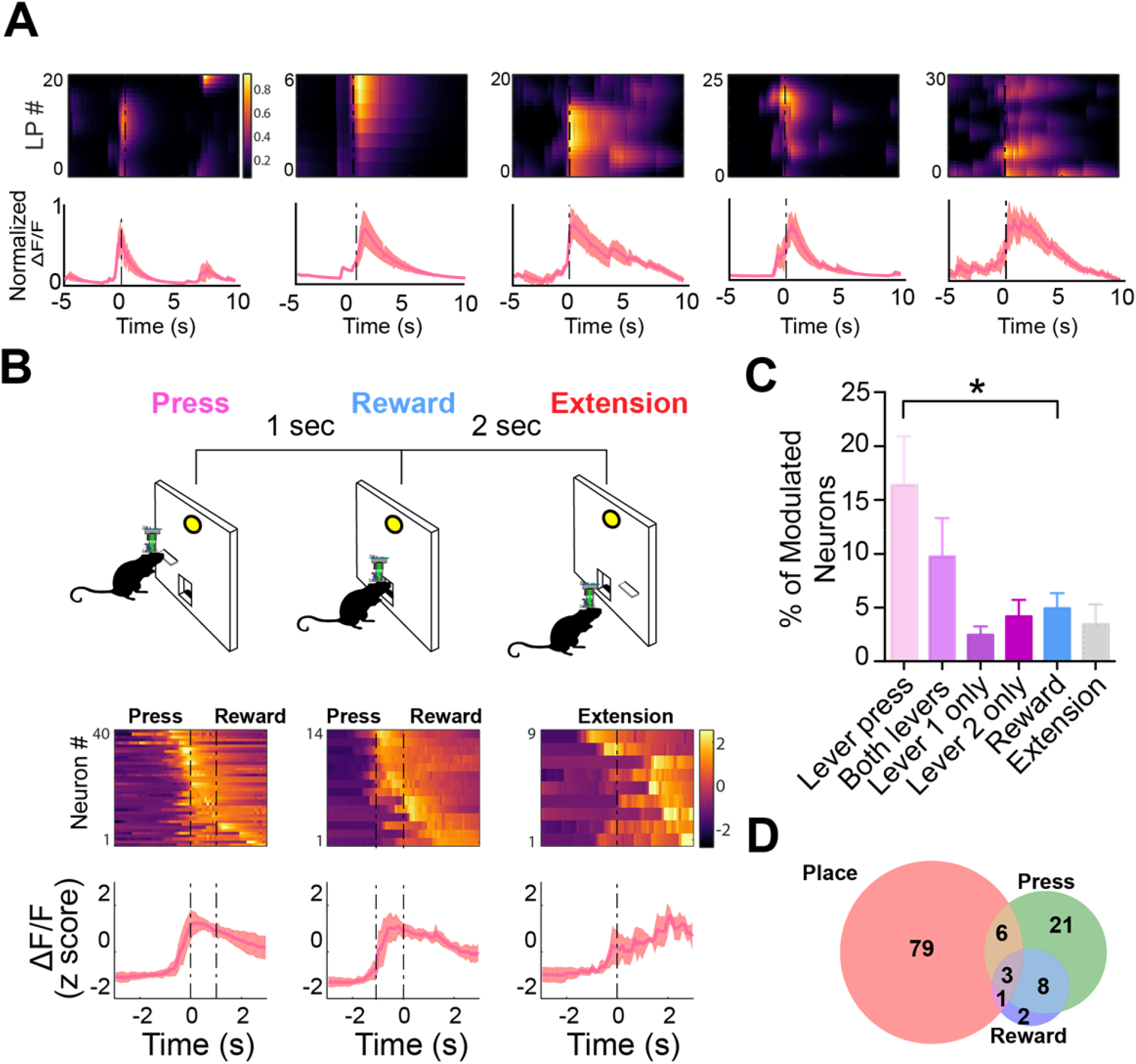
D1+ neurons in DG are significantly modulated by lever pressing, but not by passive reward delivery (see also Supplementary Figure 2). (**A**) One example neuron from each of the five calcium imaging animals, showing increased calcium transient during lever pressing. *Top*: Heat map shows normalized calcium activity aligned to a single press as a function of time during fixed-ratio (FR) 1 trials. *Bottom*: Averaged calcium activity across all presses. For the 5 different animals we recorded N=64 (Animal 1), N=62 (Animal 2), N=22 (Animal 3), N=59 (Animal 4), and N=100 (Animal 5) (**B**) *Top:* Schematic of FR1 paradigm. Animals press one of two levers that is randomly presented, which then retracts followed by pellet delivery one-second later. After two seconds, one of two levers extends again at random. *Bottom*: Peri-event heat maps and average traces of calcium activity aligned to either *all lever presses, reward*, or *lever extension*. Only neurons that are significantly modulated around each event are shown. (**C**) Percentages of modulated neurons by each event. More neurons are modulated by lever pressing than reward delivery (One-way ANOVA, *F(5,24)* = 4.1077, *p* = 0.0078; Dunnett’s multiple comparison: Lever press vs reward *p* < 0.05). (**D**) Venn-diagram displaying the number of neurons in a session modulated by spatial location (place), lever pressing, or reward delivery.

It is difficult to assess spatial activity of neurons in operant tasks, as animals do not cover the arena equally but instead preferentially occupy specific task-relevant locations (**Supplementary Figure 2**). To examine the stability of spatially related activity during operant conditioning, we used an FR5 task with two levers (**Supplementary Figure 3**). We then used the same methods as described in (Skaggs et al., 1993) to identify spatially modulated neurons, and split the sessions into periods when the left or right lever were active. This allowed us to recalculate the center of mass of our identified spatial firing fields with two different lever locations. As mice mostly stayed close to the wall where the two levers and food port were located, we limited our analysis to the x-dimension which explained most of the variance in the neural activity. We found that the place field centers were significantly different when the lever is available. While we did find that occupancy varied in this switch task, the occupancy across other tasks was consistent, suggesting that spatial modulation does not depend on the type of task (e.g. operant vs Pavlovian) (**Supplementary Figure 2**). In contrast, task-related neural activity in the DG depends on whether the task is action-contingent.

## Discussion

Together our results provide the first evidence that DG D1+ neurons play a role in operant reinforcement. Using a cell-type specific approach, we showed that activation of D1+ neurons in the DG is sufficient for self-stimulation (Gangarossa et al., 2012). Furthermore, our retrograde tracing identified that the DG receives input from the LC and not from the VTA or SNc. DG receives a TH+ projection from the LC rather than VTA, suggesting that the LC is a source of dopaminergic projections to the DG. Stimulation of the LC-DG projection was sufficient to support self-stimulation. Blockade of D1 receptors, but not noradrenergic beta receptors, attenuated the self-stimulation response of hippocampal-projecting LC neurons. These findings suggest that the LC supplies the primary dopaminergic input to the dorsal hippocampus, especially to a population of D1+ neurons in the DG, and that this dopaminergic pathway plays a critical role in operant reinforcement.

Both the hippocampus and the LC are known to be effective sites for intracranial self-stimulation (Ursin et al., 1966; Crow et al., 1972; Ritter and Stein, 1973). However, as non-selective electrical stimulation was used in classic studies, the precise circuit mechanisms underlying these observations remain unclear. In the present study, we have discovered a novel pathway for operant reinforcement that originates from LC DA neurons and targets D1+ DG neurons in the hippocampus. Self-stimulation behavior supported by stimulation of the LC-DG projections or the D1+ DG neurons appears to be different from that supported by stimulation of dopamine neurons in the VTA or SNc. Future work will have to address whether this self-stimulation effect requires D1+ neurons or if it is a property of hippocampal neurons in general.

Our calcium imaging results showed that D1+ neurons in the DG are preferentially active during goal-directed instrumental actions compared to passive reward delivery. These findings suggest that the hippocampus contributes to reinforcement of specific instrumental actions. They are broadly in agreement with recent findings on entorhinal grid cells (Butler et al., 2019) and hippocampal CA1 cells (Gauthier and Tank, 2018), as well as LC terminals in CA1 that were preferentially active near a novel reward location (Kaufman et al., 2020). It remains to be determined how the activity of DG D1+ neurons change during activation of LC neurons that project to the hippocampus, and if LC activation induces plasticity in the DG that is important for learning.

## Materials and methods

All experimental procedures were conducted in accordance with standard ethical guidelines and were approved by the Duke University Institutional Animal Care and Use Committee.

### Subjects

All behavioral data were collected from *D1-cre* mice (B6;129-Tg(Drd1-cre)120Mxu/Mmjax, Jackson Labs), and *wild-type* (C57BL/6J). Optogenetic control of D1-receptor expressing dentate hippocampal neurons was achieved with a double-floxed inverted recombinant AAV5 virus injection to express the excitatory opsin ChR2-eYFP. Viral infection in the dentate of the hippocampus was histologically verified with eYFP imaging colocalized against a D1 receptor antibody and DAPI staining. All mice were aged between 2-12 months old, housed on a 12:12 light cycle, with tests occurring in the light phase. For calcium imaging experiments, mice were put on food restriction and maintained at 90% of their initial body weights.

### Viral Constructs

CAV2-Cre was obtained from Institut de Génétique Moléculaire de Montpellier. rAAV5.EF1α.DIO.hChR2(H134R).eYFP, rAAV5.EF1α.DIO.eYFP, AAV9.hSyn.FLEX.jGCaMP7F, AAV(retro2).hSyn.EF1α.Cre.WPRE were obtained from the Duke University Vector Core. pAAV_hSyn1-SIO-stGtACR2-FusionRed was from Ofer Yizhar (Addgene viral prep # 105677-AAV1; http://n2t.net/addgene:105677; RRID:Addgene_105677).

### Pathway-specific retrograde tracing experiments

Retrograde anatomical tracing data was collected from Ai14 reporter mice (129S6-Gt(ROSA)26Sor^tm14(CAG-tdTomato)Hze^/J, Jackson labs). Ai14 reporter mice have a loxP-flanked STOP cassette that is excised in the presence of Cre to promote transcription of a CAG promoter-driven red fluorescent protein variant (tdTomato). 50 nL of either Retro2-Cre (**Figure 2**), or canine adenovirus type 2 expressing Cre recombinase (CAV2-cre, **Supplementary Figure** 1) was injected into the DG of Ai-14 reporter mice (AP: -2.0 mm relative to bregma, ML: ± 1.3 mm relative to bregma, DV: 2.0 mm from skull surface) (Soudais et al., 2001; Tervo et al., 2016).

### RNAscope

3 D1-Cre male mice were injected with the AAV9-hSyn-Flex-GCaMP7f in the DG (−2.0 & -2.2 AP; +\-1.3 ML; 1.8 DV. from bregma). 3-4 weeks after injections, mice were euthanized with CO2, and their brain was quickly harvested and frozen in OCT for future RNAscope experiments using the Advanced Cell Diagnostics kit and probes (ACD). Brains were sectioned using a cryostat at a thickness of 20 micrometers and directly mounted on superfrost slides; the slides were then stored at -80.

On the experiment day, 4% PFA in PBS was chilled at 4°C in a PFA-safe IHC container. Slides were removed from the -80 and immediately immersed in the pre-chilled PFA for 15 minutes at 4°C. After washing the slides with PBS, sections were dehydrated through a 5 minutes immersion at 50%, 70%, and 100% ethanol. Slides were then air-dried for 5 minutes at room temperature before incubating them for 30 minutes with protease IV (ACD #322336), then washed twice in PBS. Sections were then incubated with either the Drd1a probe (ACD #406491), the negative control probe (ACD #320871), or the positive control probe (ACD #320881) for 2 hours at 40°C. Next, slides were washed twice using the wash buffer (ACD #310091) and then incubated for 30 minutes with Amp1, 15 minutes with Amp2, 30 minutes with Amp3, and finally 15 minutes with Amp4 Alt B-FL (ACD #320851). After the two washes, sections were then incubated with 5% Neutral Goat Serum (NGS) in 0.2% TBST for 1 hour at RT in the dark, then incubated for 1 hour with a primary antibody against GFP (1:1000; Millipore, AB16901) and for 2 hours with a secondary Alexa-fluorophore (488) conjugated antibodies (Invitrogen). Slides were mounted in Vectashield with DAPI (Vector Laboratories, CA) and a minimum of 5 images per mouse were acquired on an Olympus Fluoview confocal microscopy using a 60X oil immersion objective.

Images were then processed using FIJI (https://imagej.net/Fiji/Download), GFP+ cells were identified and saved as individual ROIs. Using the GFP+ cells, a mask was created to identify the presence of puncta (probe positive signals) within each ROI using the puncta analyzer plugin. The percentage of GFP+ cells having puncta and the average number of puncta per GFP+ cell was calculated in all three conditions. (**Figure 6**)

### Histology and immunohistochemistry

Mice were anesthetized and transcardially perfused with 0.1M phosphate buffered saline (PBS) followed by 4% paraformaldehyde (PFA) in order to confirm viral expression as well as optic fiber and GRIN lens placement. To confirm placement, brains were stored in 4% PFA with 30% sucrose for 72 hrs. Tissue was then post-fixed for 24 hours in 30% sucrose before cryostat sectioning (Leica CM1850) at 60 µm. Fiber and lens implantation sites were then verified. To confirm eYFP expression in LC and DG neurons, sections were rinsed in 0.1M PBS for 20 min before being placed in a PBS-based blocking solution. The solution contained 5% goat serum and 0.1% Triton X-100 and was allowed to sit at room temperature for 1 hr. Sections were then incubated with a primary antibody (polyclonal rabbit anti-TH 1:500 dilution, ThermoFisher, catalog no. P21962; polyclonal chicken anti-EGFP, 1:500 dilution, Abcam, catalog no. ab13970) in blocking solution overnight at 4 °C. Sections were then rinsed in PBS for 20 min before being placed in a blocking solution with secondary antibody used to visualize *TH* neurons in the LC (goat anti-rabbit Alexa Fluor 594, 1:1000 dilution, Abcam, catalog no. ab150080; goat anti-chicken Alexa Fluor 488, 1:1000 dilution, Life Technologies, catalog no. A11039) for 1 hr at room temperature. Sections were mounted and immediately coverslipped with Fluoromount G with DAPI medium (Electron Microscopy Sciences; catalog no. 17984-24). Placement was validated using an Axio Imager.V16 upright microscope (Zeiss) and fluorescent images were acquired and stitched using a Z780 inverted microscope (Zeiss).

### Co-localization analysis with tracing

In order to characterize projections to the dentate gyrus we injected 50 nL of AAV(retro2).hSyn.EF1α.Cre.WPRE into each hemisphere of the DG of Ai14 mice (four mice x two hemispheres, N=8; 2 females and 2 males). We then processed the slices and acquired images as described above. We opened the raw images taken from the Axio Imager V16 upright microscope (Zeiss) in Fiji to quantify the number of cells from a single coronal brain slice using 8-bit confocal images. A threshold was set to identify the neuronal cell bodies. The function “fill holes” was then used to remove possible empty space within the selected cells. After converting the image to mask, we ran the “Analyze Particle” plug-in in Fiji to count the cells in each image. Using the Analyze Particle function, the masks taken were then counted to determine the number of co-localizing cells using the “Colocalization Threshold” plug-in in Fiji.

### Optogenetic experiments

Mice were anesthetized with 2.0 to 2.5% isoflurane mixed with 1.0 L/min of oxygen for surgical procedures and placed into a stereotactic frame (David Kopf Instruments, Tujunga, CA). Meloxicam (2 mg/kg) and bupivacaine (0.20 mL) were administered prior to incision. To optogenetically interrogate D1+ neurons in the hippocampus, adult *D1-cre* mice were randomly assigned to D1::ChR2(DG) (*n =* 8, 5 males, 3 females, 8-10 weeks old) or D1::eYFP(DG) groups (*n =* 5, 2 males, 3 females, 8-10 weeks old). Craniotomies were made bilaterally above the hippocampus and AAV5-DIO-ChR2 was microinjected into the dentate gyrus through a pulled glass pipette (200 nL each hemisphere at 1nL/s, AP: 2.0 mm relative to bregma, ML: ±

1.3 mm relative to bregma, DV: 2.0 mm from skull surface) using a microinjector (Nanoject 3000, Drummond Scientific). Optic fibers (SFLC230-10; 200 um core, 0.35 aperture, Ø1.25 mm, 6.4 mm Long SS Ferrule for MM Fiber, Ø231 µm Bore Size) were then implanted bilaterally above the dentate gyri. For pathway-specific experiments, *wild type* mice were used to selectively target LC-Hipp (*n =* 6, 3 males, 3 females, 8-10 weeks old) neurons by bilaterally injecting AAV(retro2).hSyn.EF1α.Cre.WPRE into the dentate gyrus (150 nL each hemisphere) in parallel with a Cre-dependent ChR2 virus injection into the locus coeruleus (AP: -5.45 mm relative to bregma, ML: ± 1.10 mm relative to bregma, DV: 3.65 mm from skull surface) before optic fiber placement (AP: -5.45 mm relative to bregma, ML: ± 1.10 mm relative to bregma, DV: 3.50 mm from skull surface), or eYFP (*n=6*, 3 males, 3 females). To further examine this pathway we used D1-Cre animals with the pathway specific approaches discussed above (*n =* 5, 3 males, 2 females, 8-10 weeks old) with simultaneous inhibition of D1+ neurons in the DG. To inhibit these neurons we injected Cre-dependent soma-targeted *Guillardia theta* anion-conducting channelrhodopsin (AAV1-hSyn-SIO-stGtACR2-FusionRed). An additionally 5 mice (4 male, 1 female) were used to isolate the neurotransmitters released from the LC into the DG. LC and DG surgeries were the same as pathway specific manipulations, with the addition of cannulas (P1 technologies. AP: -2.00 mm relative to bregma, ML: ± 1.8 mm relative to bregma, DV: -1.4 mm, at a 10 degree angle). All optic fibers were secured in place with dental acrylic adhered to skull screws. Mice were group housed and allowed to recover for one week before experimentation.

### Operant self-stimulation

Standard operant boxes (Model ENV-007, MED Associates, Inc., Albans, VT) were housed in light and sound attenuating cubicles (Model ENV-019, MED Associates, Albans, VT). Each box is equipped with two levers. A Windows XP based computer system running MED-PC Version IV Research Control & Data Acquisition System software (Med Associates, St. Albans, VT) is used to control the experimental equipment and record the data.

For self-stimulation experiments a single lever was inserted at the start of the session. For each lever press animals received 500 ms of stimulation at 20 Hz (15 ms pulse width, 5 mW power). Animals were trained with 30 minute sessions. Animals were tested for 32 consecutive days, and received 8 sessions of FR1, FR3, and FR5, and then 8 extinction sessions. To test for extinction, the lever was inserted but no stimulation was delivered for lever presses.

### Drug injections

Even low doses of DA antagonists are known to reduce self-stimulation, but low doses of NE antagonists have no effect on self-stimulation behavior (Rolls et al., 1974). In the present study, we selected doses that minimized effects on movement or arousal. The same mice used above were retrained after extinction (3 FR1 sessions) to press a lever for stimulation of LC neurons that projected to the dentate gyrus. They were then tested with DA or NE antagonists (N=8; 3 males, 5 females). Mice alternated between testing days where they received an intraperitoneal injections (SCH23390 (Tocris) at 0.1 mg/kg and 0.2 mg/kg, propranolol (Sigma Aldrich) at 3 mg/kg or 6 mg/kg, or vehicle (phosphate buffered saline), and training days with no injections. Each mouse had one testing day for each dose and drug combination (5 total), and the order of the injections was determined pseudo randomly. Injections were given 30 minutes prior to the start of the session.

For local drug infusions with chronically implanted cannulae, we trained naive animals (N=5) for 8 days on self-stimulation and used the same experimental testing protocol for infusions of SCH23390 (1.8 mM or 3.6 m) or propranolol (10.5 mM 21 mM). The dose is determined based on previous work{Takeuchi, 2016 #8385}. The drugs were infused at a rate of 0.0005mL.

### Calcium imaging experiments

In order to target the dentate gyrus, AAV9-syn-FLEX-jGCamp7f was injected (50 nL) in four penetrations (A.P -2.0, M.L. 1.3, D.V. -2.2, -2.1, -2.0, -1.9) of *D1-Cre* mice (*n =* 5, 5 males, 8-10 weeks old). A gradient index (GRIN) lens (Inscopix: 1 mm x 4 mm, 1.8 mm DV) was then implanted over the injection site. Viral expression was checked three weeks post-injection, and a base plate was secured to the skull with dental cement. A UCLA miniscope was used to assess in vivo activity of D1+ neurons in the DG in freely moving animals. Images were collected from the miniscope using Bonsai (Lopes et al., 2015). This allowed for simultaneous collection of calcium data with behavioral videos (Logitech c920). Calcium traces were then motion corrected https://github.com/flatironinstitute/CaImAn) and extracted using constrained nonnegative matrix factorization for micro endoscopic data (https://github.com/zhoupc/CNMF_E).The extracted traces were then analyzed using custom MATLAB scripts. Imaging sessions lasted 8-10 minutes.

### Fixed-ratio training with food reward (calcium imaging)

The behavioral tests used for calcium imaging were performed while the mice were food deprived to 85% of their free-feeding weight. There were 6 tasks used, with three days of testing for each task, for a total of 18 testing days. Each imaging session lasted approximately 10 minutes. Five mice were trained on a fixed ratio (FR) 1 task where they received a pellet for pressing a lever. Subsequently, animals were moved to an FR3, and then FR5.

To examine the interaction of spatial location and lever pressing we used an *FR5 switch task*. Where a single lever (**Supplementary Figure 3**) into an operant chamber, and 5 responses resulted in a pellet. Following 5 pellets (25 presses), the first lever retracted, and a second lever is inserted. The mouse has to move to the other side of the food cup and press the other lever to earn a food reward.

In order to dissociate reward delivery and action production we used an FR1 schedule of reinforcement where the reward was delayed one second after the press (*Two-lever FR1 delay task)*. We used two levers in this task. One lever is inserted on a given trial. If pressed the lever would retract, and a food pellet is delivered 1 s later. After a 2 s inter-trial-interval, one of the two levers was randomly inserted.

Following the fixed ratio testing, mice received a reward at fixed intervals (20s), preceded by 1 second of white noise (*Non-contingent reward)*. The animals were not required to press a lever to receive the reward.

### Analysis

To assess significant increases in calcium activity, for each neuron we made a peri-event time histogram (PETH) between -3 s and 3 seconds aligned to relevant events(da Silva et al., 2018). Event times were considered between -500ms and +500ms around the relevant event. Baseline activity was considered -3000 to -1000 ms prior to the event. If the calcium activity was 99% above baseline for 3 consecutive 100 ms bins then there was considered to be a significant increase in calcium activity. For tasks that involved two levers we calculated the percent of neurons that were significantly modulated by either lever, and then excluded these neurons from our analysis of modulation by a specific lever. We identified spatially tuned cells by computing the spatial information contained in the calcium transients, compared with shuffled data(Skaggs et al., 1993). The data arena was split into a 15 by 15 bins (2×2 cm each), and neurons were required to be active in active in at least three bins to be considered spatially modulated.

## Acknowledgments

This work was supported by NIH grants DA040701 and MH112883 to H.H.Y.

## Supplementary Figures

**Supplementary Figure 1:**
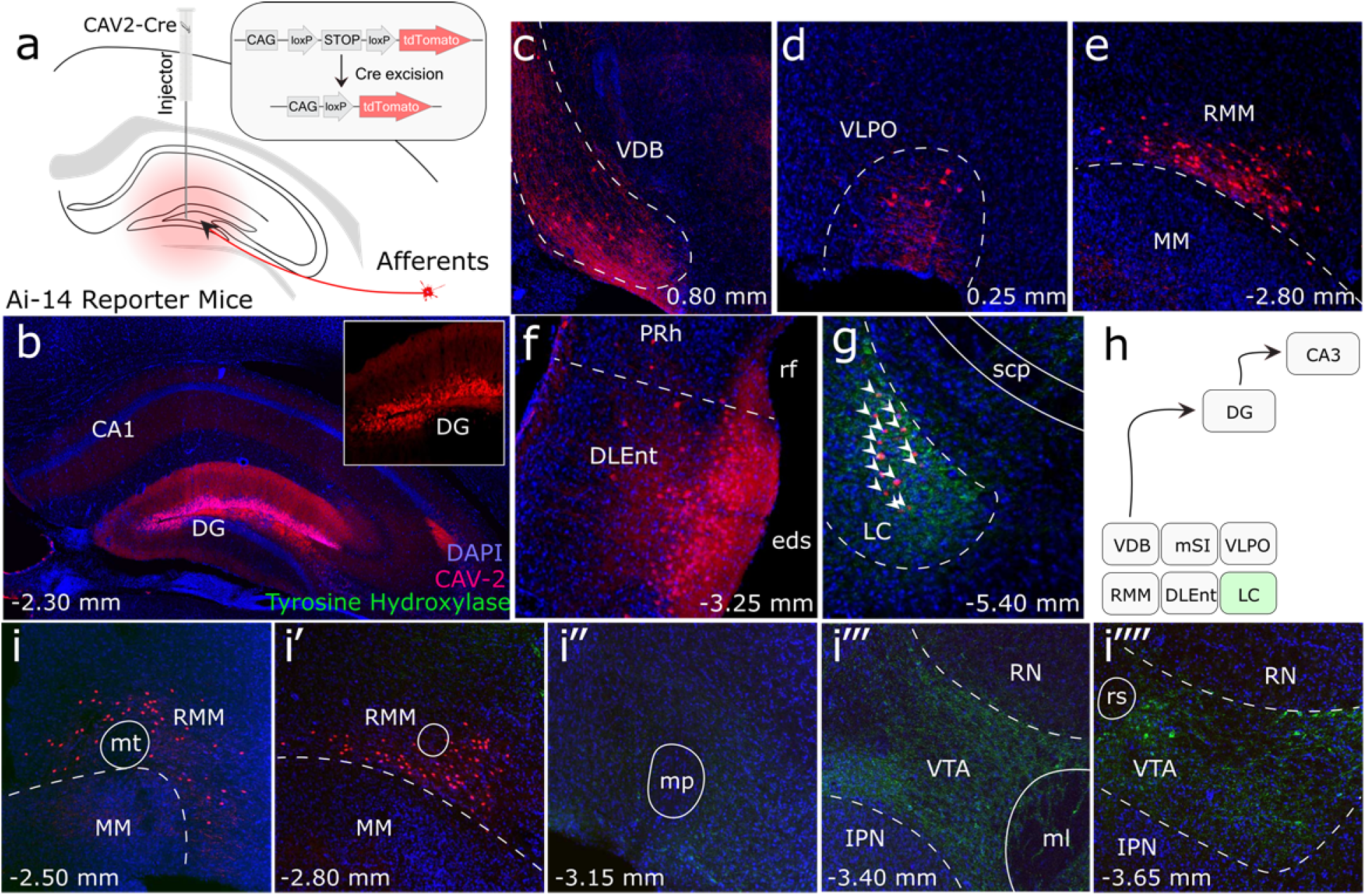
Retrograde tracing shows Locus Coeruleus (LC) projections, but not Ventral Tegmental Area (VTA) projections to the dorsal dentate gyrus. (**a**) Schematic of Cav2-Cre injection into the dentate gyrus of the hippocampus of Ai-14 reporter mice. (**b**) Injection site of the Cav2 showing the Cre-positive neurons. (**c-f**) Retrograde labeling of neurons in canonical brain regions that project to the hippocampus. Vertical diagonal band (VDB), Ventral lateral preoptic area (VLPO), retro mammillary bodies (RMM), dorsal lateral entorhinal cortex (DLEnt). (**g**) Retrograde labeling of neurons in the LC, colocalized with tyrosine hydroxylase (TH). (**h**) Schematic summarizing all brain regions that project to the DG. Only LC is TH+ (**i - i’’’’**) Anterior to posterior series of brain sections, demonstrating that retrograde labeling ends outside of the VTA.

**Supplementary Figure 2:**
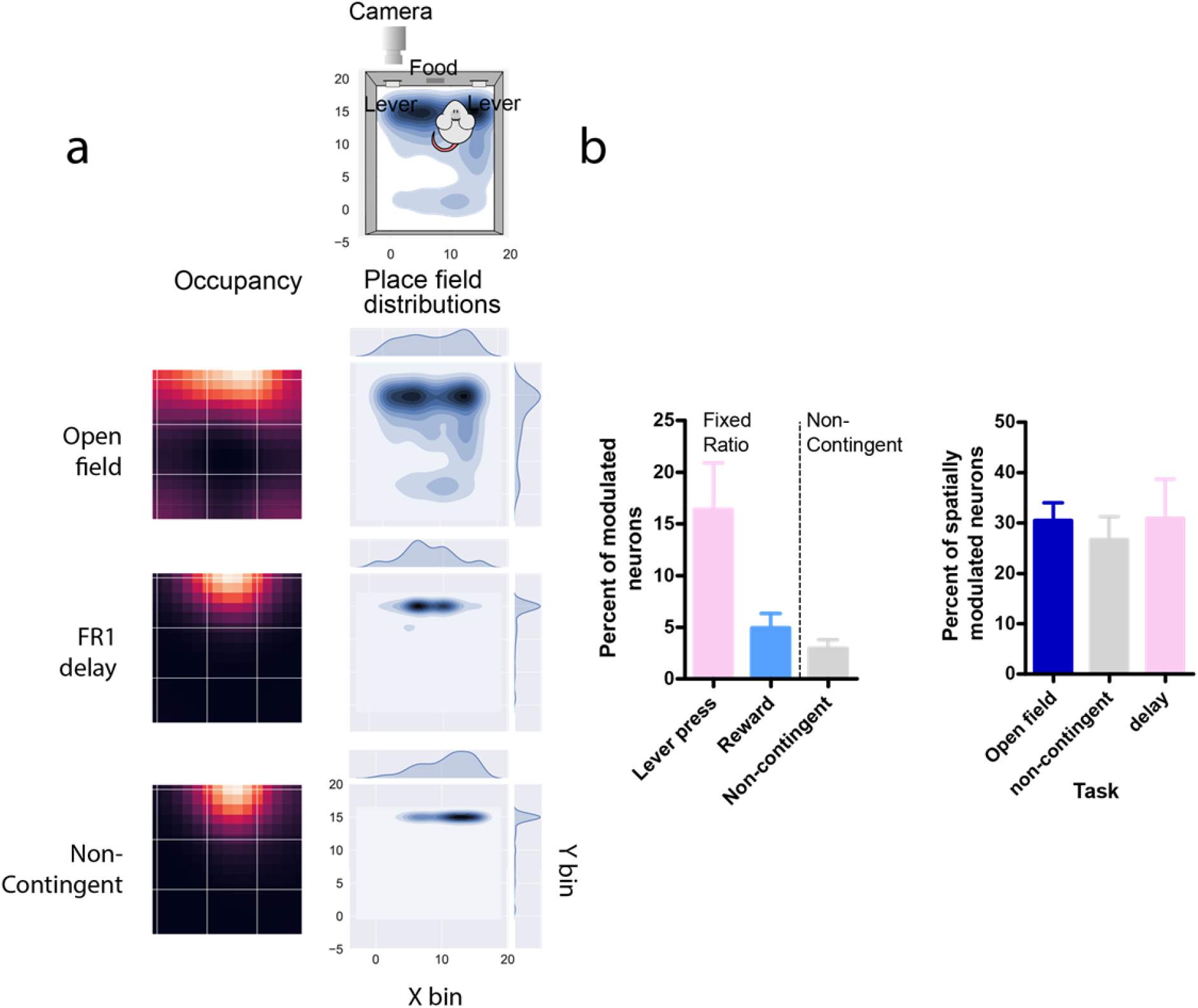
Comparison of neural activity in D1+ DG neurons during different behavioral tasks. (**a**) Top down views of the arena. Left column, occupancy of the animal during the task. Right column, kernel density estimation plots and corresponding distributions showing the center of mass of spatially modulated cells during different tasks. The occupancy and place field distribution largely overlap. (**b**) Left, percent of modulated neurons for specific events during either an FR1 delay task, or a non-contingent reward task. We found higher number of neurons active during lever pressing compared to reward delivery in the same task, or reward delivery in a non-contingent task. Right, The proportion of spatially modulated neurons during an open-field baseline session, non-contingent reward task, or an FR1-delay task.

**Supplementary Figure 3:**
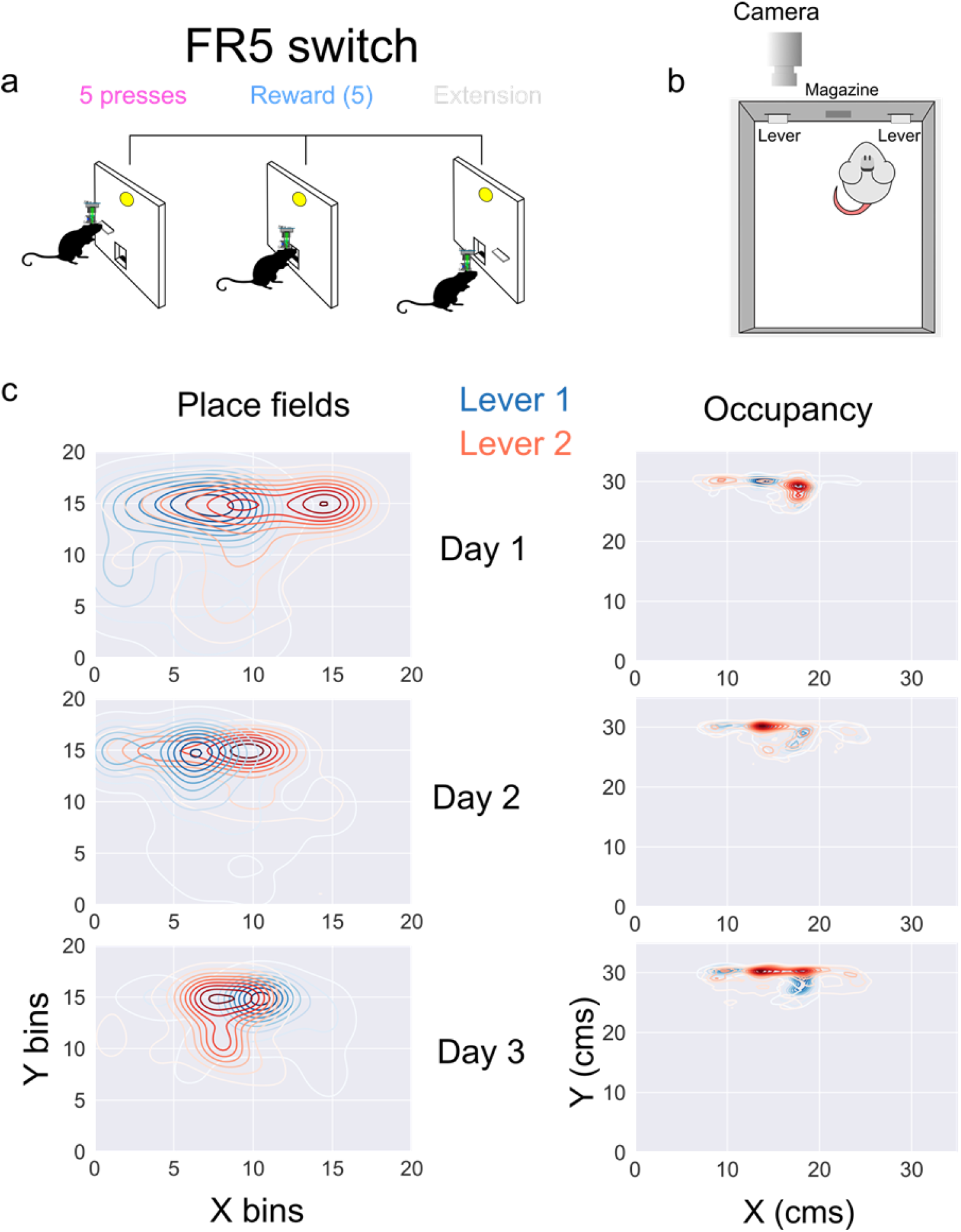
Comparison of spatially modulated D1+ DG neurons while pressing two different levers. (**a**) A schematic of the task, where 5 presses results in reward delivery. Once animals receive 5 rewards (25 presses) the lever retracts and a second lever extends. Animals were tested for three days, and data was pooled across days here. (**b**) Top down schematic of the experimental setup with two levers. (**c**) *Left*, place field centers for lever 1 (blue) and lever 2 (red), here the data is binned into 2 cm bins, and kernel density estimations are plotted (paired samples t-test: [Day1, L1 vs L2] N-51, p-3.44e-0.7, [Day2, L1 vs L2] N-72, p-0.00186, [Day3, L1 vs L2] N-73, p-0.000304. Holm-Bonferroni correction for multiple comparisons = 0.0166). *Right*, occupancy when lever 1 or lever 2 is extended.

## Notes

### Competing Interest Statement

The authors have declared no competing interest.

## References

Adcock RA, Thangavel A, Whitfield-Gabrieli S, Knutson B, Gabrieli JD (2006) Reward-motivated learning: mesolimbic activation precedes memory formation. Neuron 50:507–517.

Björklund A, Dunnett SB (2007) Dopamine neuron systems in the brain: an update. Trends in neurosciences 30:194–202.

Butler WN, Hardcastle K, Giocomo LM (2019) Remembered reward locations restructure entorhinal spatial maps. Science 363:1447–1452.

Crow TJ, Spear P, Arbuthnott G (1972) Intracranial self-stimulation with electrodes in the region of the locus coeruleus. Brain Research 36:275–287.

da Silva JA, Tecuapetla F, Paixão V, Costa RM (2018) Dopamine neuron activity before action initiation gates and invigorates future movements. Nature 554:244.

Eldridge LL, Knowlton BJ, Furmanski CS, Bookheimer SY, Engel SA (2000) Remembering episodes: a selective role for the hippocampus during retrieval. Nature neuroscience 3:1149–1152.

Gangarossa G, Longueville S, De Bundel D, Perroy J, Hervé D, Girault JA, Valjent E (2012) Characterization of dopamine D1 and D2 receptor-expressing neurons in the mouse hippocampus. Hippocampus 22:2199–2207.

Gauthier JL, Tank DW (2018) A dedicated population for reward coding in the hippocampus. Neuron 99:179-193. e177.

Ikemoto S (2007) Dopamine reward circuitry: two projection systems from the ventral midbrain to the nucleus accumbens-olfactory tubercle complex. Brain Res Rev 56:27–78.

Kaufman AM, Geiller T, Losonczy A (2020) A Role for the Locus Coeruleus in Hippocampal CA1 Place Cell Reorganization during Spatial Reward Learning. Neuron.

Kempadoo KA, Mosharov EV, Choi SJ, Sulzer D, Kandel ER (2016) Dopamine release from the locus coeruleus to the dorsal hippocampus promotes spatial learning and memory. Proceedings of the National Academy of Sciences 113:14835–14840.

Kravitz AV, Owen SF, Kreitzer AC (2013) Optogenetic identification of striatal projection neuron subtypes during< i> in vivo</i> recordings. Brain research 1511:21–32.

Lopes G, Bonacchi N, Frazao J, Neto JP, Atallah BV, Soares S, Moreira L, Matias S, Itskov PM, Correia PA, Medina RE, Calcaterra L, Dreosti E, Paton JJ, Kampff AR (2015) Bonsai: an event-based framework for processing and controlling data streams. Frontiers in neuroinformatics 9:7.

Milner B, Squire LR, Kandel ER (1998) Cognitive neuroscience and the study of memory. Neuron 20:445–468.

Mishkin M, Malamut, B., & Bachevalier, J. (1984) Memories and habits: Two neural systems. In: Neurobiology of learning and memory (Lynch G, J. L. McGaugh, N. Weinberger, ed), pp 65-77. New York: Guilford Press.

Morris R, Anderson E, Lynch Ga, Baudry M (1986) Selective impairment of learning and blockade of long-term potentiation by an N-methyl-D-aspartate receptor antagonist, AP5. Nature 319:774–776.

Ritter S, Stein L (1973) Self-stimulation of noradrenergic cell group (A6) in locus coeruleus of rats. Journal of Comparative and Physiological Psychology 85:443.

Rolls E, Kelly P, Shaw S (1974) Noradrenaline, dopamine, and brain-stimulation reward. Pharmacology Biochemistry and Behavior 2:735–740.

Rossi MA, Sukharnikova T, Hayrapetyan VY, Yang L, Yin HH (2013) Operant Self-Stimulation of Dopamine Neurons in the Substantia Nigra. PloS one 8:e65799.

Schultz W, Dayan P, Montague PR (1997) A neural substrate of prediction and reward. Science 275:1593–1599.

Schwarz LA, Luo L (2015) Organization of the locus coeruleus-norepinephrine system. Current Biology 25:R1051–R1056.

Skaggs WE, McNaughton BL, Gothard KM (1993) An information-theoretic approach to deciphering the hippocampal code. In: Advances in neural information processing systems, pp 1030–1037.

Soudais C, Laplace-Builhe C, Kissa K, Kremer EJ (2001) Preferential transduction of neurons by canine adenovirus vectors and their efficient retrograde transport in vivo. The FASEB Journal 15:1–23.

Takeuchi T, Duszkiewicz AJ, Sonneborn A, Spooner PA, Yamasaki M, Watanabe M, Smith CC, Fernández G, Deisseroth K, Greene RW (2016) Locus coeruleus and dopaminergic consolidation of everyday memory. Nature 537:357–362.

Tervo DG, Hwang BY, Viswanathan S, Gaj T, Lavzin M, Ritola KD, Lindo S, Michael S, Kuleshova E, Ojala D, Huang CC, Gerfen CR, Schiller J, Dudman JT, Hantman AW, Looger LL, Schaffer DV, Karpova AY (2016) A Designer AAV Variant Permits Efficient Retrograde Access to Projection Neurons. Neuron 92:372–382.

Tsai HC, Zhang F, Adamantidis A, Stuber GD, Bonci A, de Lecea L, Deisseroth K (2009) Phasic firing in dopaminergic neurons is sufficient for behavioral conditioning. Science 324:1080–1084.

Ursin R, Ursin H, Olds J (1966) Self-stimulation of hippocampus in rats. Journal of Comparative and Physiological Psychology 61:353.

Wise RA (2004) Dopamine, learning and motivation. Nature reviews Neuroscience 5:483–494.

Yin HH, Ostlund SB, Knowlton BJ, Balleine BW (2005) The role of the dorsomedial striatum in instrumental conditioning. The European journal of neuroscience 22:513–523.

Yttri EA, Dudman JT (2016) Opponent and bidirectional control of movement velocity in the basal ganglia. Nature 533:402–406.

